# On the Periodic Gain of the Ribosome Flow Model

**DOI:** 10.1101/507988

**Authors:** Mahdiar Sadeghi, M. Ali Al-Radhawi, Michael Margaliot, Eduardo Sontag

**Affiliations:** Department of Electrical and Computer Engineering, Northeastern University; Department of Electrical Engineering-Systsems, Tel Aviv University

## Abstract

We consider a compartmental model for ribosome flow during RNA translation called the Ribosome Flow Model (RFM). This model includes a set of positive transition rates that control the flow from every site to the consecutive site. It has been shown that when these rates are time-varying and jointly *T*-periodic every solution of the RFM converges to a unique periodic solution with period *T*. In other words, the RFM entrains to the periodic excitation. In particular’ the protein production rate converges to a unique *T*-periodic pattern. From a biological point of view, one may argue that the average of the periodic production rate, and not the instantaneous rate, is the relevant quantity. Here, we study a problem that can be roughly stated as: can periodic rates yield a higher average production rate than constant rates? We rigorously formulate this question and show via simulations, and rigorous analysis in one simple case, that the answer is no.

## Introduction

Transcription and translation are the two major steps of gene expression, that is, the transformation of the information encoded in the DNA into proteins. During translation complex molecular machines called ribosomes traverse the mRNA molecule, “read” it codon by codon, and generate the corresponding chain of amino-acids.

New imaging techniques [1, 2, 3, 4] and empirical approaches [5, 6, 7, 8, 9, 10] for studying gene expression provide unprecedented amounts of data on the dynamics of translation. This increases the need for mathematical and computational models for ribosome flow that can integrate, explain and make predictions based on this data (see the reviews [11, 12, 13]). Mechanistic models are particularly important in biotechnology and synthetic biology, as they allow to predict the effect of various manipulations of the biological machinery [14, 15].

The Ribosome Flow Model (RFM) is a deterministic model for ribosome flow [16]. It can be derived via a dynamic mean-field approximation of a fundamental model from statistical physics called the Totally Asymmetric Simple Exclusion Process (TASEP) [11, 17]. In TASEP particles hop randomly along a chain of ordered sites. Totally asymmetric means that the flow is unidirectional, and simple exclusion means that a particle can only hop into a free site. This models the fact that two particles cannot be in the same place at the same time. Note that this generates an indirect coupling between the particles. In particular, if a particle is delayed at a site for a long time then the particles behind it cannot move forward and thus a “traffic jam” may evolve.

The RFM is a compartmental model with n sites. The state-variable *x_i_*(*t*), *i* = 1,…, *n*, describes the density of particles at site *i* at time *t*. This is normalized so that *x_i_*(*t*) = 0 [*x_i_*(*t*) = 1] means that site *i* is completely empty [full] at time *t*.

The dynamics is described by *n* first-order ODEs:

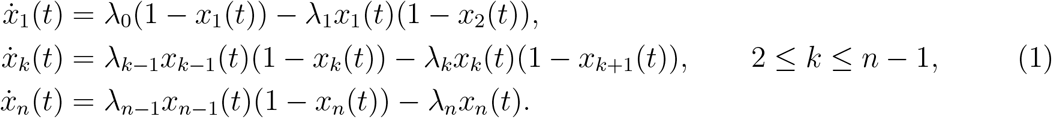

Here *λ_í_* > 0 is a parameter that describes the translation rate from site *i* to site *i* + 1, with *λ*_0_ [*λ_n_*] called the entry [exit] rate. Eq. (1) can be explained as follows. The flow from site *k* to site *k* + 1 is given by *λ_k_x_k_*(1 − *x*_*k*+1_), i.e. it increases when site *k* becomes fuller and decreases when site *k* + 1 becomes fuller. This is a “soft” version of simple exclusion. The production rate at time *t* is the rate of ribosomes exiting site *n*, that is, *R*(*t*) := *λ_n_x_n_*(*t*).

Note that if some λ_*k*_ is small then the flow from site *k* to site *k* + 1 will be small, so site *k* fills up. Consequently, the flow λ_*k*−1_ *x*_*k*−1_(1 − *x*_*k*_) from site *k* − 1 to site *k* will become small and then site *k* − 1 fills up. In this way, a traffic jam may evolve behind a “bottleneck” site. The implications of such traffic jams in various biological transportation processes is recently attracting considerable interest (see, e.g. [18, 19]).

It has been shown in [20] that there exists a unique *e* = *e*(λ_0_,…, λ_*n*_) ∈ (0,1)^*n*^ such that any solution of the RFM emanating from the unit cube converges to *e*. Thus, the system is globally asymptotically stable. In particular, the production rate converges to

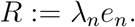

Ref. [21] derived a spectral representation for the steady-state density *e* and production rate *R*. Given the RFM, define the (*n* + 2) × (*n* + 2) tridiagonal matrix

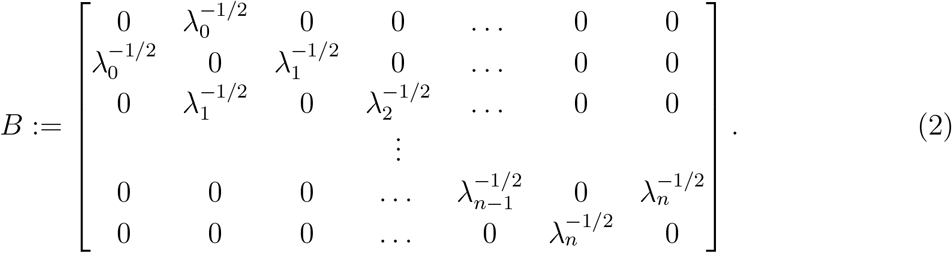

Note that *B* is (componentwise) nonnegative and irreducible. Let 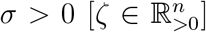 the Perron root [Perron vector] of *B*. Then

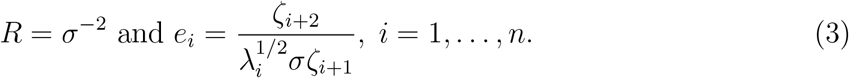

Note that it follows from this that for any *c* > 0 we have

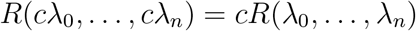

that is, the steady-state production rate is positively homogeneous of degree one.

### Example 1.

*Consider the RFM with all the rates equal to one. Then B is a tridiagonal Toeplitz matrix and it is well-known (see e.g. [22]) that its eigenvalues are 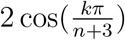, k = 1,…, n + 2, so the Perron root is 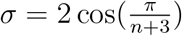. The corresponding Perron vector is*

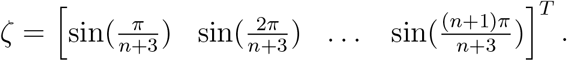

*Thus, in this case* (3) *gives*

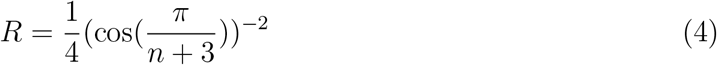

*and*

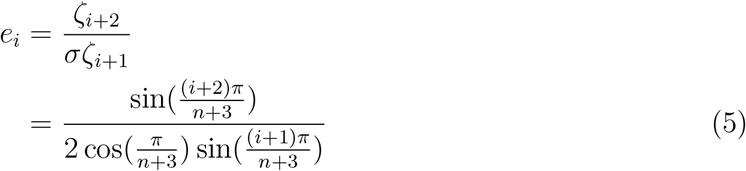

*for all i = 1,…,n. For example, in the one-dimensional case, i.e. 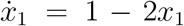 the equilibrium point is e* = 1/2, *so R* =1/2, *whereas* (4) *yields*

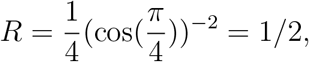

*and* (5) *gives* 
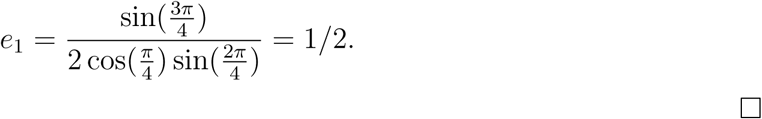

Biological organisms are exposed to periodic excitations like the 24h solar day and the periodic cell-cycle division process. Proper functioning often requires entrainment to such excitations i.e. internal processes must operate in a periodic pattern with the same period as the excitation. An example is the sleep-wake cycle that entrains to the 24h day.

Ref. [23] studied the RFM with positive time-varying rates that are jointly *T*-periodic, and proved that every state-variable converges to a periodic solution with period *T*. In other words, the RFM entrains. The proof is based on the fact that the RFM is an (almost)contractive system [24]. However, this provides no information on the attractive periodic solution (except for its period).

Since any set of jointly periodic rates induces a periodic solution, a natural question is: can periodic rates yield a higher production rate than constant rates? In this paper, we formulate this question rigorously and then address it via simulations and rigorous analysis in one simple case. The remainder of this paper is organized as follows. The next section defines the periodic gain of the RFM. Section 3 describes some simulation results for the general RFM. Section 4 considers the problem of finding the maximal periodic gain as an optimal control problem. We then turn to a particular simple case, namely, a one-dimensional RFM, and prove a result on the periodic gain using the PMP.

## 2 Problem Formulation

For any *T*-periodic function *f*, with *T* > 0, let 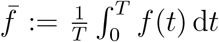, that is, the average of *f* over a period. Pick a set of rates λ_*i*_(*t*), *i* = 1,…,*n*, that are jointly *T*-periodic (note that a constant rate is *T*-periodic for any *T*). Recall that this induces a unique *T*-periodic trajectory *γ*(*t*) of the RFM and thus a unique *T*-periodic production rate *R_T_*(*t*) := λ_*n*_(*t*)*γ_n_*(*t*). The average production rate is thus 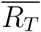. Consider an RFM with constant rates 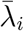, *i* = 1, …, *n*. Recall that every trajectory converges to a unique steady-state e and thus to a production rate 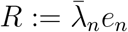.

The question we are interested in is: what is the relation between 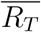 and *R*? Note that this is a “fair” comparison as we replace every time-varying rate by its average value.

We call 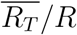 the *periodic gain* of the RFM (for the given set of *T*-periodic rates). One can argue that the average production rate over a period, rather than the instantaneous value, is the biologically relevant quantity. Then a periodic gain larger than one implies that we can “do better” using periodic rates.

To gain a wider perspective, consider the case of a SISO asymptotically stable linear system with input [output] *u*(*t*) [*y*(*t*)] and transfer function *G*(*s*). Suppose that *u*(*t*) = *a* + *b* sin(*ωt*). Note that this is *T*-periodic with *T* = 2*π/ω*, and that 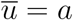. It is well-known that the output converges to the *T*-periodic function *y_T_*(*t*) := |*G*(0)|*a* + |*G*(*jω*)|*b* sin(*ωt* + ∠*G*(*jω*)), where 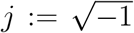, so 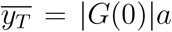. On the other-hand, if we replace *u*(*t*) by 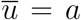 then the output converges to |*G*(0)|*a*. Thus, the periodic gain for this input is one and by superposition it is one for any periodic input.

Of course, for nonlinear systems, like the RFM, the periodic gain may be different than one. The next example demonstrates this.

### Example 2.

*Consider the scalar system*

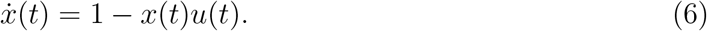

*For u*(*t*) = 1 + (1/2) cos(*t*) the solution is

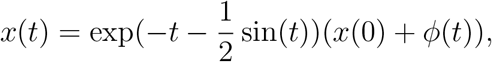

*where* 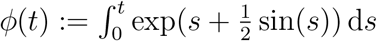. *Thus*,

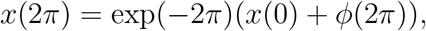

*We now determine an initial condition x*(0) = *c for which the solution is 2π-periodic, that is*,

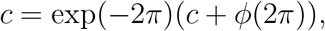

*so*

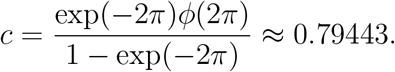

*Thus, the periodic solution is* 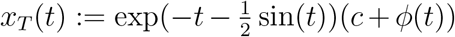. *It is not difficult to show that this solution is attractive. A calculation yields 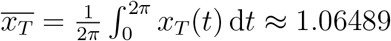*.

*On the other-hand, for 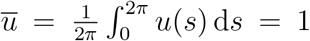, the solution of* (6) *converges to the steady-state* 1 *and thus the periodic gain is* ≈ 1.06489.

## 3 Simulations

We consider the RFM where every rate is a sum of m harmonic functions with random coefficients. More precisely, we generated a matrix of random entries *P* ∈ ℝ^(*n*+1)×(2*m*)^ and then set

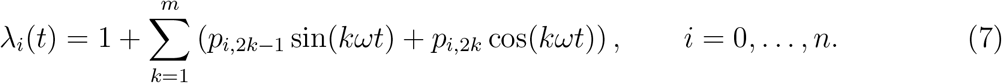

Note that this guarantees that the λ_*i*_’s are jointly *T*-periodic for *T* = 2*π/ω* and that 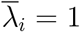 for all *i*. The entries of *P* are generated randomly with a uniform distribution over [−1/(2*m*), 1/(2*m*)], so that λ_*i*_j(*t*) ≥ 0 for all *i* and all *t*.

We first simulated the RFM with *n* =1. Since 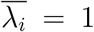 for *i* = 0,1 we know from Example 1 that *R* = 1/2. Fig. 1 depicts a histogram of the average steady-state flow 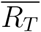 for *m* = 3 and 10,000 simulations. It may be seen that this is always smaller than 1/2. Thus, the constant rates yield the maximal production rate.

**Figure 1:**
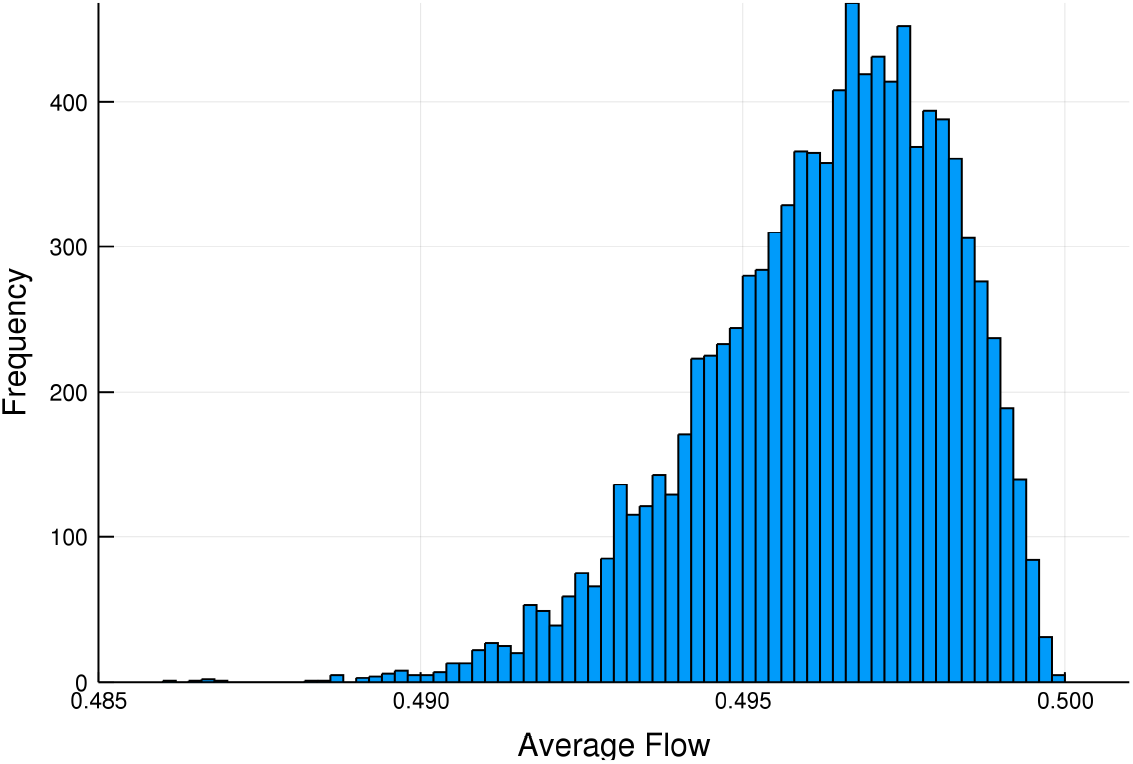
A histogram of 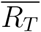 values in the one-dimensional RFM with periodic λ_0_(*t*) and λ_1_(*t*)

Consider now an RFM with *n* = 4 and constant rates set to one. By (4),

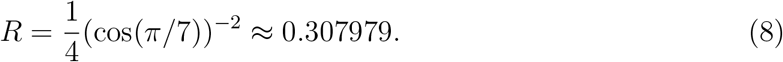

Fig. 2 depicts a histogram representation of 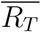 calculated over 10,000 simulations of an RFM with periodic rates λ_0_(*t*),…, λ_4_(*t*). It may be seen that 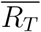 is smaller than the value in (8), so again the constant rates are those that maximize the production rate.

**Figure 2:**
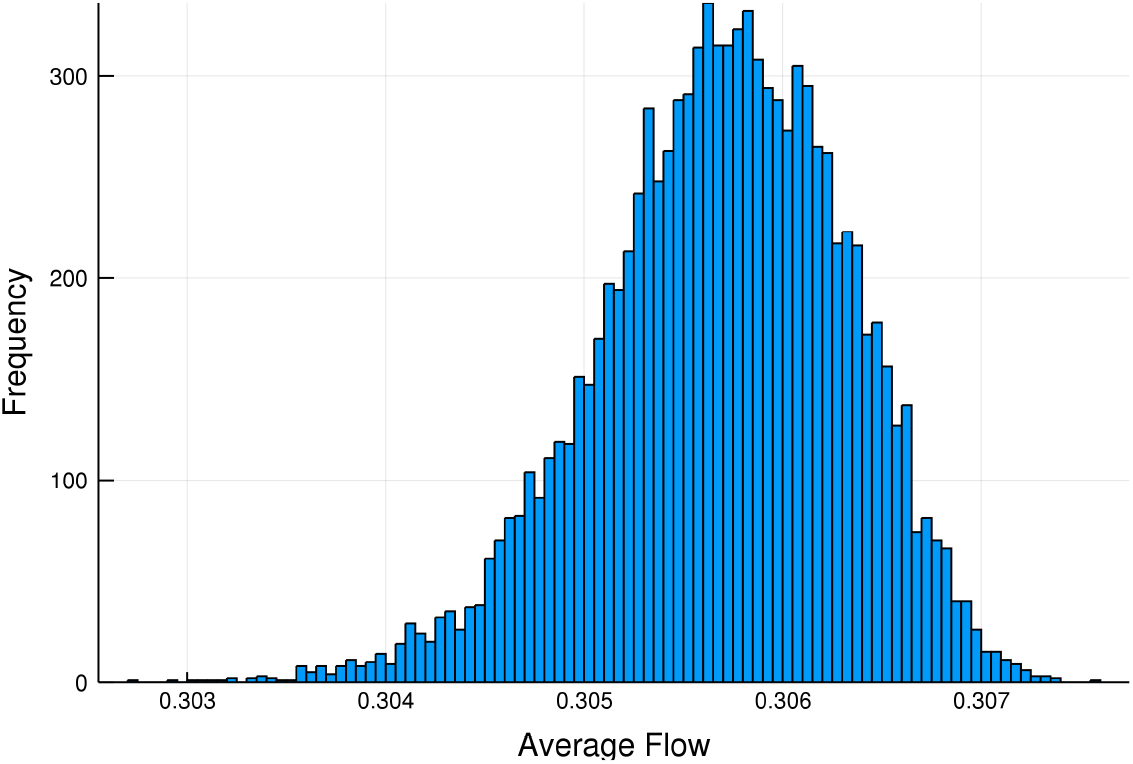
A histogram of 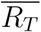 values in an RFM with *n* = 4 and periodic rates λ_*i*_(*t*), *i* = 0,…, 4.

Our next goal is to pose the problem of determining the RFM periodic gain as an optimal control problem.

## 4 Optimal periodic control

We augment the equations of the RFM with an additional state-variable as follows

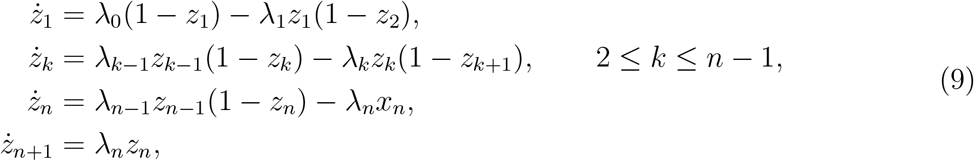

with *z*_*n*+1_(0) = 0. Thus, 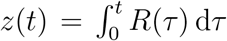. Pick *T* > 0 and *a*_1_,…, *a_n_* > 0, and let *S_a,T_* denote the set of measurable functions λ*j*(*t*), *i* = 0,…, *n*, satisfying the following conditions for all *i* = 0, …, *n*,

1. λ_*i*_(*t*) ≥ 0 for almost all *t* ≥ 0;
2. λ_*i*_(*t*) = λ_*i*_(*t* + *T*) for almost all *t* ≥ 0; and
3. 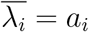.

We can apply the spectral representation to determine the steady-state production rate *R* when λ_*i*_(*t*) ≡ *a*_*i*_ for all i. Thus, determining the periodic gain is equivalent to solving the following optimization problem.

### Problem 1.

*Maximize* 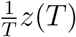 *over the set of admissible rates S_a,T_ and with the boundary condition z*_*i*_(0) = *z*_*i*_(*T*) ∈ (0,1) for *i* = 1,…, *n*.

The last condition guarantees that we are maximizing the production rate along the (unique) periodic trajectory.

## 5 Optimal Control and Pontryagin Principle

### 5.1 Optimal Control Formulation

Consider the following single state system:

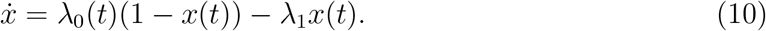

Let λ_0_(*t*) : ℝ_≥0_ → ℝ_>0_ be a measurable signal with a period *T* > 0, i.e, λ_0_(*t*) = λ_0_(*t* + *T*) > 0, and let λ_1_ > 0 be a constant.

Let *φ*(*t*; *x*(0), λ_0_(*t*)) be the solution of (10) for a given inflow rate and an initial condition (which exists and is unique by Carathéodory conditions [25]). Since the system is contractive, then the solution of (10) entrains to the periodic input, i.e, it converges to a periodic trajectory with a period *T* [23]. Let *x*(*t*) : φ*(*t*; λ_0_) be the limit trajectory.

The problem in the last section can be understood as maximizing the average flow, i.e *y*(*t*) = λ_1_*x*(*t*), over the set of inputs with a given average.

In order to write this as an optimal control problem, we consider optimizing the following cost functional over the compact interval [0, *T*]:

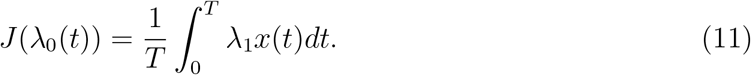

In order to enforce the periodicity condition, we impose the constraint *x*(0) = *x*(*T*). The set of admissible controls is the set of measurable signals λ_0_(*t*) over the interval [0, *T*] with a given average 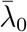 and which satisfy λ_0_(*t*) ∈ [0, *L*] for some upper bound *L* > 0. The periodic extension of a signal can be defined as λ_0_(*t*) = λ(t − *mT*), where 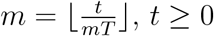.

Hence, we write the extended system as:

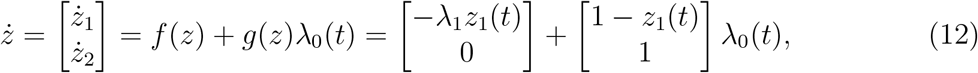

where *z*_1_(*t*) := *x*(*t*), and with the following boundary conditions:

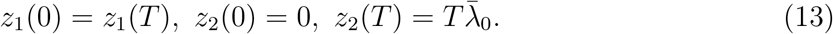

*For a given L* > 0, *the optimal control problem can be stated as follow:*

#### Problem 1.

*Find* λ_0_(*t*) : [0, *T*] → [0, *L*] *that maximizes the cost functional* (11) *subject to the ODE* (12) *and the boundary conditions* (13).

Note that Problem 1 is a nonlinear optimal control problem since it has a bilinear term containing the state and control.

The first main result is the following:

#### Theorem 1.

*The signal that solves Problem 1 with* 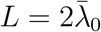 *is:*

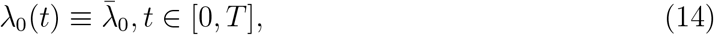

*and corresponding the optimal cost is:*

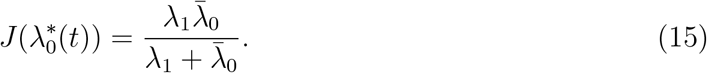

Hence, the throughput of the system can not be improved by using periodic time-varying inflows. The proof of Theorem 1 is given below.

### 5.2 Pontryagin’s Maximum Principle with periodic inputs

The Pontryagin Maximum Principle (PMP) is a well-known necessary condition for the optimality of control signals [26],[27],[28]. However, Problem 1 has a non-standard boundary condition *z*_1_(0) = *z*_2_(T). Nevertheless, we can use a modified transversality condition to state the PMP. First, we need to define the Hamiltonian associated with the system as:

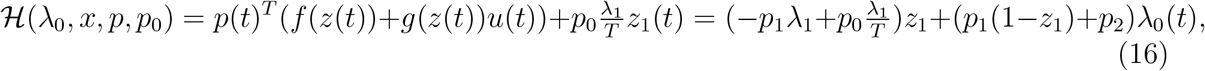

where *p*(*t*) ∈ ℝ^2^ is the costate, and *p*_0_ ≥ 0 is called the abnormal multiplier.

For Problem 1, the maximum principle can be state as:

#### Proposition 2

(The Maximum Principle). *Let* 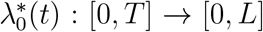 *be an optimal control for Problem 1. Let z*\* : [0, *T*] → [0, 1] × *R be the corresponding optimal state trajectory. Let* 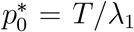. *Then, there exists a function p\** : [0, *T*] → ℝ^2^ *with p\**(*t*) ≠ 0 *for all t* ∈ [0, *T*], *and satisfying:*

1. The solution z*(*t*) *and p\**(*t*) *satisfy:*

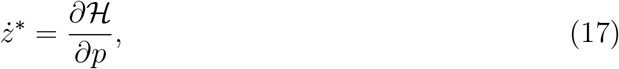

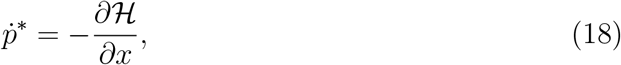

*where* 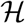 *is given by* (16) *with the boundary conditions* (13).
2. *The control* 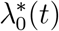 *is optimal for all t* ∈ [0, *T*] *while fixing z*,p*, i.e*,

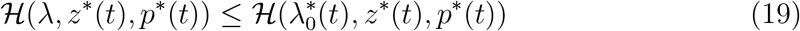

*for all t* ∈ [0, *T*] *and* λ ∈ [0, *L*].
3. *The costate variable satisfies the following transversality condition:*

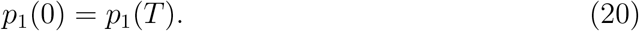

*Proof*. We need to prove the transversality condition and that the abnormal multiplier is non-zero

For the transversality condition, we utilize a result from [27] that allows for arbitrary initial sets. Let *S* ⊂ ℝ^4^ be an *initial set*, and have the following constraint:

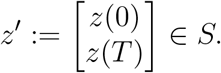

Then transversality condition for the PMP can be stated as:

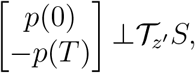

where 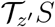 is the tangent space of *S* at *z′*.

In our case, 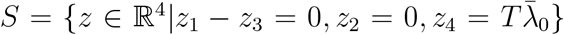. Hence, *T_z_S* = span{[1, 0, 1, 0]^*T*^}. Therefore, it is necessary that 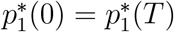.

Next, we show that abnormal multiplier can not vanish. For the sake of contradiction, assume *p*_0_ = 0. Hence, (18) implies that 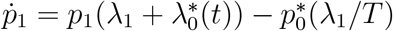. Therefore,:

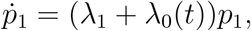

which is separable. Hence, integrating we get 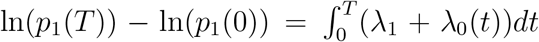. Using the transversality condition derived above this implies that 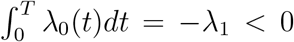 which contradicts the fact that λ_0_(*t*) > 0. Hence, 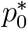 can not vanish. Assume now that a pair 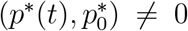 exists with 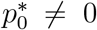 that satisfies the PMP. Then, 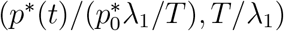 also satisfies the PMP. Hence, we can state the PMP with 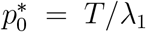 without loss of generality.

## 6 Optimality of the constant rate

### 6.1 Characterization of the optimal rate

The control input λ_0_(*t*) appears linearly in the the Hamiltonian (16), and also *p*_2_(*t*) ≡ *p*_2_(0) which follows from (18) since 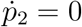. Hence, we can define *the switching function* as:

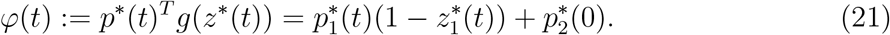

We have the following lemma:

#### Lemma 3.

*Let* 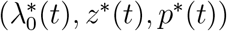 *be an optimal trajectory, then if φ*(*t*) ≠ 0, *then:*

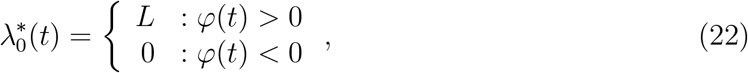

*i.e*, λ_0_(*t*) is a bang-bang control when the switching function does not vanish.

*Proof*. Let φ(*t*) > 0, and let 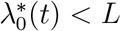. However,

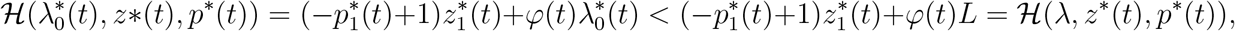

which violates condition 3 in Proposition 2. Hence, 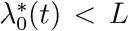 is not optimal. The same argument can be applied when φ(*t*) < 0.

The switching function can also vanish. In that case the control signal is called *singular*. We provide a characterization of the form of the singular control. First we need some notation. Assume that 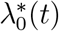 is an optimal control and let φ be the associated switching function. Let *E*^+^ = {*t* ∈ [0, *T*]|φ(*t*) > 0}, *E*^−^ = {*t E* [0, *T*]|φ(*t*) < 0}, and *E*_0_ = {t ∈ [0, *T*]|φ(*t*) = 0}. Note that all three sets are measurable. We state the characterization next.

#### Lemma 4.

*Let* 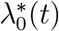 *be an optimal control, and let E*^+^, *E*^−^, *E*_0_ *be as defined above such that μ*(*E*_0_) > 0, *where μ denotes the Lebesgue measure. Then:*

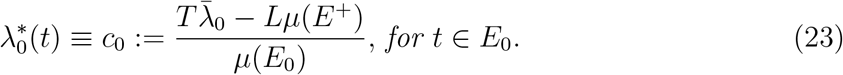

*and* 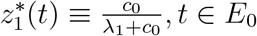.

*Proof*. Let *E* ⊂ *E*_0_ be the set of accumulation points of *E*_0_. Note *μ*(*E*) = *μ*(*E*_0_), since *E*_0_ – *E* is the set of isolated points of *E* which is countable, and hence has measure zero. Then, φ(*t*) = 0 for *t* ∈ *E* implies that *d^n^*/*dt^n^*(*φ*(*t*)) = 0, *t* ∈ *E* for all positive integers *n*.

Hence, we calculate 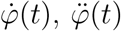 below. Lie brackets are usually utilized for such calculation.

Using φ(*t*) = *p*_1_(1 – *z*_1_) + *p*_2_(0), we can write:

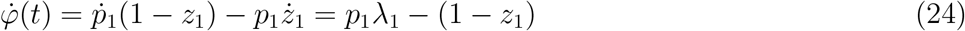

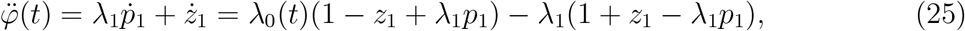

In order to find the form of the singular trajectory on *E*, we find that 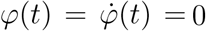 implies that 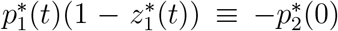, and 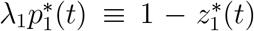. Hence, we find that 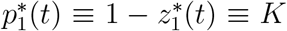, where *K* is a constant. Since *z*(*t*) ∈ (0,1), then *K* ∈ (0,1). Hence, 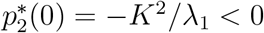.

Substituting 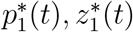 in 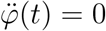 we get that:

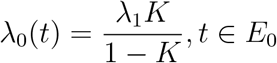

which is a constant. Using the fact that 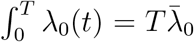, and noting that λ_0_(*t*) ∈ {0, *c*_0_, *L*} we can write 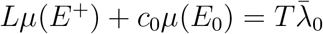, and hence (23) follows (i.e, 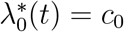), and 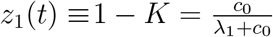.

Lemma 1 and 4 above lead to the following characterization of optimal control signals:

#### Proposition 5.

*Let* 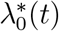 *be a control input that solves Problem 1, then* λ_0_(*t*) ∈ {0, *c*_0_, *L*}, *for some c*_0_ ∈ (0, *L*).

#### Remark 1.

*If μ*(*E*_0_) = 0 *then the control signal is said to be* fully bang-bang, *while if μ*(*E*_0_) = *T then the control signal is* fully singular. *Otherwise, the control signal is said to be a patching of* bang-bang arcs *and* singular arcs. *Hence, Theorem 1 asserts that the control is fully-singular. In that case* 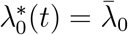 *and* 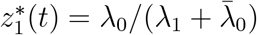.

#### Remark 2.

*The singular control satisfies the necessary conditions such as the Legendre-Clebsch condition [27]. However, we will not include a proof as we will be showing that it is indeed optimal within the class of three-level signals*.

### 6.2 Any patching of bang-bang with singular arcs is suboptimal

Proposition 5 shows that an optimal control takes up to three values only. Hence, in order to prove Theorem 1 we need to show that the cost functional with a fully singular control achieves better than any other signals in that class. We parametrize, the class of signals for a given average 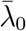 as follows:

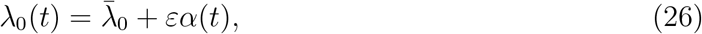

where *α*(*t*) is measurable such that *α*(*t*) ∈ { –1,*c*, 1}, where |*c*| < 1 and 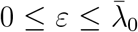. It also satisfies 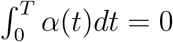. For every choice of λ_0_(*t*) we let *x*(*t*) denote the solution of (10) that satisfies *x*(0) = *x*(*T*).

We will show that 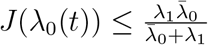 for any choice *ε* and *α*(*t*) above.

First, we study the case when *α*(*t*) has a *finite* number of switchings, which we define next. A set *E* ⊂ [0, *T*] is said to be *elementary* if it can be written as a *finite* union of open, closed, or half-open intervals. Let *E*^+^, *E*^−^, *E*_0_ be defined similar to the previous subsection, i.e, we let *E*^+^ = {*t*|*α*(*t*) = 1}, *E*^−^ = {*t*|*α*(*t*) = −1}, *E*_0_ = {*t*|*α*(*t*) = *c*}. Then *α*(*t*) is said to have a finite number of switchings if *E*^+^, *E*^−^, *E*_0_ are elementary sets.

We are ready to state the next proposition:

#### Proposition 6.

*Let* λ_0_(*t*) *be given as in* (26) *such that α*(*t*) *has a finite number of switchings*.

*Then*,

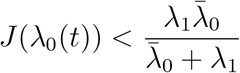

*for ε* > 0, *μ*(*E*_0_) < *T*.

The proof will be presented after some preliminaries.

First, we derive an identity for the scalar linear system 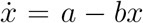, where *a*, *b* > 0 are constants. This linear system has a unique steady state at 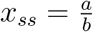 which is globally stable. We state the following lemma:

#### Lemma 7.

*Let t*_0_, *t*_1_ > 0 *be the initial and final time. Consider the initial value problem* 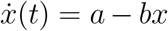 *with x*(*t*_0_) = *x*_0_. *Let x*(*t*) *be solution of the problem, and let x*_1_ = *x*(*t*_1_). *Then:*

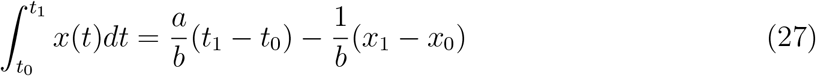

*Proof*. Since the scalar linear system is separable, we can write: 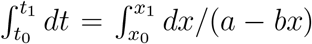.

Hence,

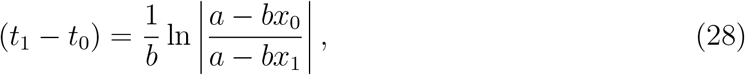

which implies

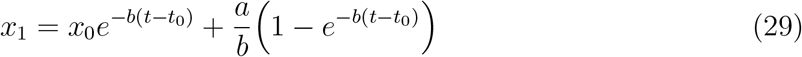

Next, we show that the integral of *x*(*t*) over time can transform to integration over occupancy:

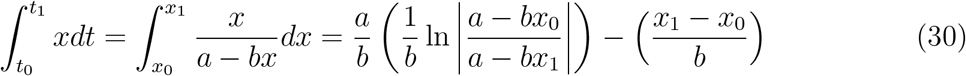

Using (28) we get the required formula (27).

Going back to our problem, note that the system (10) with an input (26) is a *switched linear system* which switches between three linear systems. Fig. 3 illustrates this switched system with two lines in 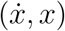 coordinates, where:

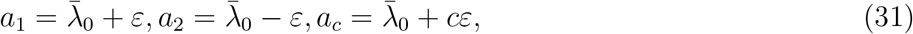

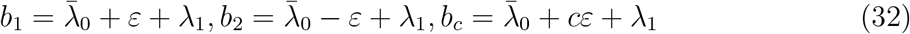

**Figure 3:**
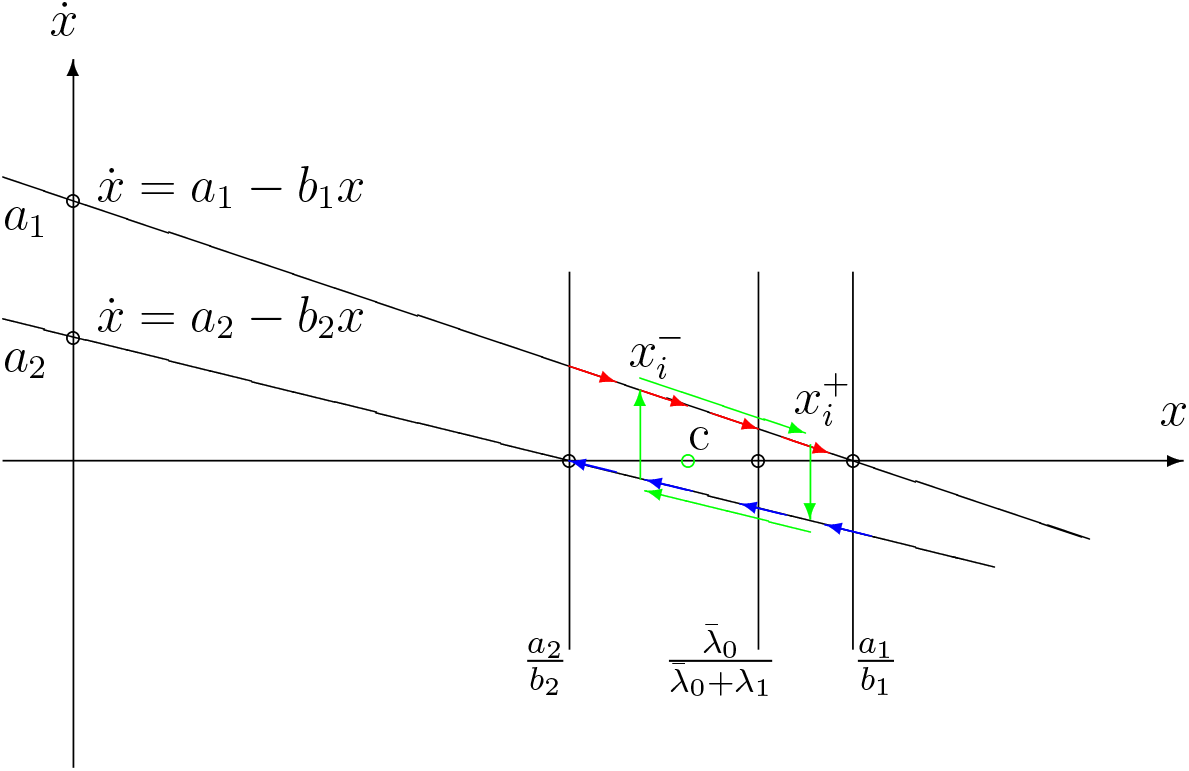
A trajectory of the switched system in the 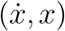-coordinates.

The upper line is when *α*(*t*) = 1, and the lower line is when *α*(*t*) = −1. Note that there is no line for the case *α*(*t*) = *c* since the characterization in Lemma 4 implies that *x*(*t*) is constant on a singular arc, and hence 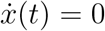. Hence, when the control is singular, the system 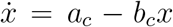 is at steady-state, and the trajectory stays at a single point in 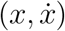-coordinates.

We need the following lemma to characterize the behaviour of the system for a given *α*(*t*) with finite switchings:

#### Lemma 8.

*Let* λ_0_(*t*) *given with a finite number of switchings, and let x*(*t*) *the solution of* (10) *with x*(0) = *x*(*T*). *Let t^c^* = *μ*(*E*_0_). *Then there exists a positive integer n, and* 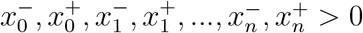 *such that:*

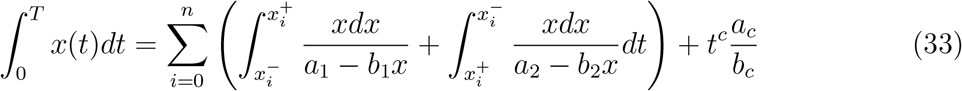

*Proof*. The trajectory starts from *x*(0) and returns to *x*(0) at *t* = *T*, which forms a grand loop. (note that the time spent when *α*(*t*) = *c* is part of the loop). Since *α*(*t*) can have multiple swtichings, it can have multiple loops. We observe the following: if the trajectory transverses from 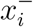 to 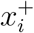 on the upper line, then it must transverse back from 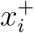 to 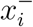 on the lower line. Hence any trajectory can be partitioned into a finite number of trajectories. The ith trajectory consists of two segments: a segment on the upper line (from 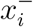 to 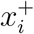) matched with a segment on the lower line (from 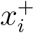 to 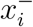). Note that the two segments are not necessarily consecutive in time, and the partitioning is not necessarily unique. Using the same transformation used in (30) and accounting for the total time spent in the singular arc, we can write the integral as in (33).

Before completing the proof of Proposition 6, we state following lemma:

#### Lemma 9.

*The following inequality holds:*

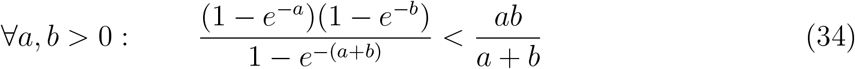

*Proof*. Let 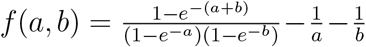. The inequality is proved if we show that *f*(*a*, *b*) > 0 for *a*, *b* > 0. Note that lim_(*a*, *b*)→(0,0),*a*,*b*>0_ *f*(*a*, *b*) = 0.

Differentiating with respect to *a*:

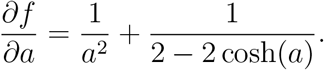

Using the Taylor series of cosh(*a*) we can find that 2cosh(*a*) − 2 > *a*^2^ for *a* > 0. Hence *∂f*/*∂a* > 0. Similarly, *∂f*/*∂b* > 0. Hence, *f* increases in all directions in the positive quarter which proves the lemma.

We complete the proof of Proposition 6 below

*Proof of Proposition 6*. Let *t^c^* = *μ*(*E*_0_), *t*^+^ = *μ*(*E*^+^), *t*^−^ = *μ*(*E*^−^). Using the fact that total time is *T*, and that 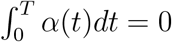. Then *t*^+^ and *t*^−^ can be written as a function of *t^c^* and *T* (35):

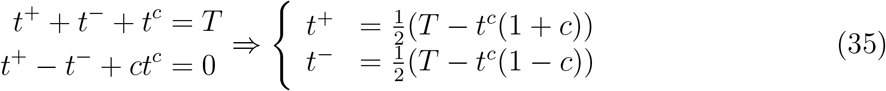

Now consider the decomposition stipulated in Lemma 8, with 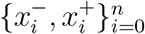. Let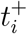 be the time spent on the upper segment from 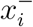 to 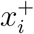, and let 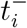 be the time spent on the lower segment from 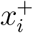 to 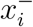.

Using (33) and (28), we can write:

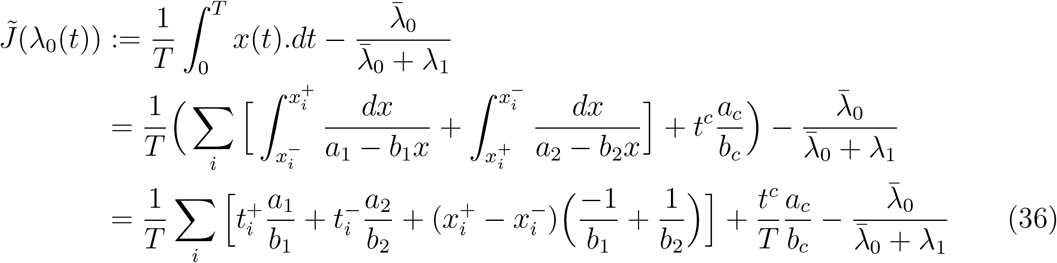

We proceed to rewrite 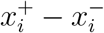 using equation (29) as follows:

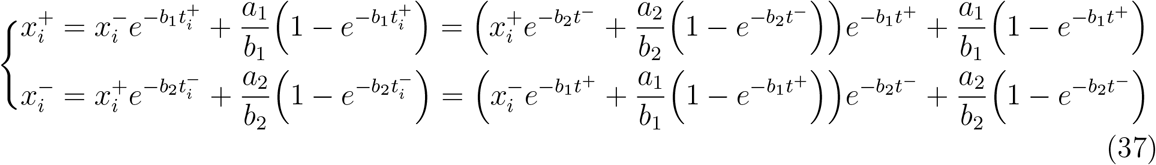

Hence,

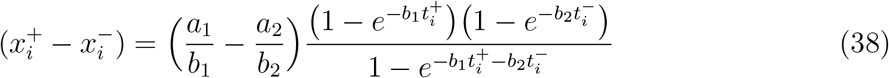

Using Lemma (9) and (38) we get the following upper bound:

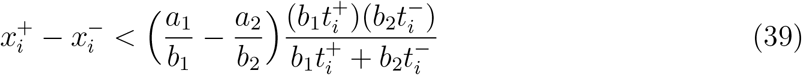

Rewriting the equation (36):

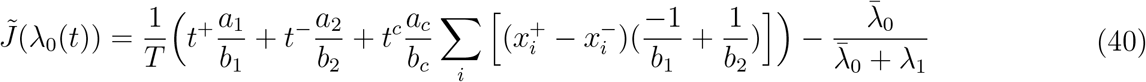

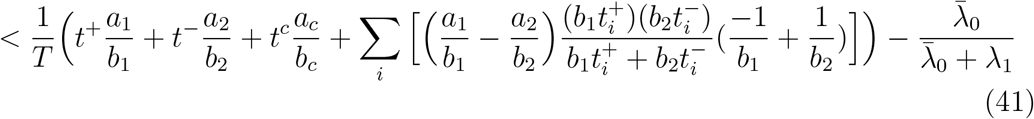

Let 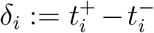. Note that it can be either positive or negative. Note that 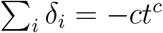. Define 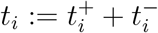, and hence 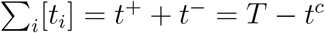. Using the definitions, we get:

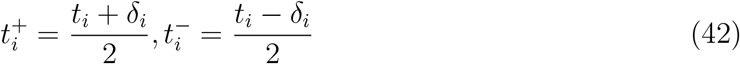

Hence, we can write (41) as follows:

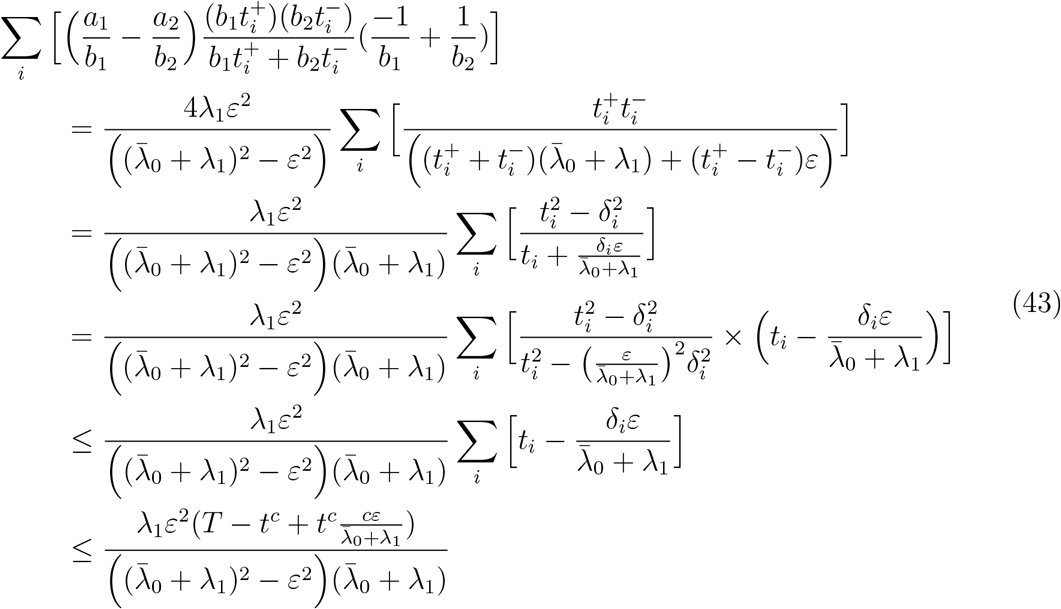

Substituting (43) in the inequality (36) and substituting the values of all the variables, we get that (44) simplifies to:

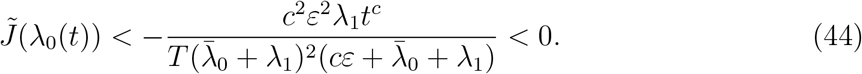

In order to prove Theorem 1, we need to consider an arbitrary measurable signal *α*(*t*). For that purpose, use the following characterization of measurable sets:

#### Lemma 10.

*[29] Let E* ⊂ [0, *T*]. *Then E is measurable if and only if for every ε* > 0 *there exists an elementary set B_ε_* ⊂ [0, *T*] *such that μ*(*E*∆*B_ε_*) < *ε*, *where* ∆ *is the symmetric difference of sets*.

We improve on the lemma above, by the following:

#### Lemma 11.

*Let E* ⊂ [0, *T*]. *Then E is measurable if and only if for every ε* > 0 *there exists an elementary set B_ε_* ⊂ [0,*T*] *with μ*(*B_ε_*) = *μ*(*E*) *such that μ*(*E*∆*B_ε_*) < *ε*, *where* ∆ *is the symmetric difference of sets*.

*Proof*. Sufficiency is clear. For necessity, let *B*_*ε*/2_ be the elementary set given by Lemma 10. We can modify the intervals contained in *B*_*ε*/2_ by up to *ε*/2 to get *B_ε_* with *μ*(*B_ε_*) = *μ*(*E*).

We generalize Proposition 6 as follows:

#### Proposition 12.

*Let* λ_0_(*t*) *be given as in* (26) *such that α*(*t*) *is measurable. Then*,

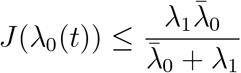

*Proof*. Let *E*^0^, *E*^+^, *E*^−^ be defined as before. Using Lemma 11, let 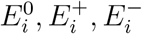 be elementary sets such that 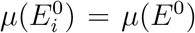 and 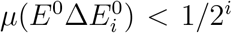, and similarly for 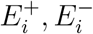. We have also 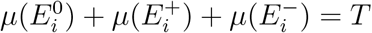.

Let *α_i_*(*t*) be defined as follows:

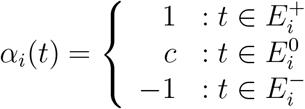

Then *α_i_*(*t*) are elementary simple functions, and we have *α_i_*(*t*) → *α*(*t*) for all *t*. For each *i*, *x_i_*(*t*) is the solution of the corresponding differential equation which has a known form. Hence, the proposition follows using Proposition 6 and Lebesgue’s bounded convergence theorem [29].

Theorem 1 follows from Propositions 5 and 12.

